# Mast cells promote pathology and susceptibility in tuberculosis

**DOI:** 10.1101/2024.09.04.611333

**Authors:** Ananya Gupta, Vibha Taneja, Javier Rangel Moreno, Nilofer Naqvi, Abhimanyu, Yun Tao, Mushtaq Ahmed, Kuldeep S. Chauhan, Daniela Trejo-Ponce de León, Gustavo Ramírez-Martínez, Luis Jiménez-Alvarez, Cesar Luna-Rivero, Joaquin Zuniga, Deepak Kaushal, Shabaana A. Khader

## Abstract

Tuberculosis (TB), caused by the bacterium *Mycobacterium tuberculosis (Mtb*), infects approximately one-fourth of the world’s population. We reported an increased accumulation of mast cells (MCs) in the lungs of macaques with active pulmonary TB (PTB), compared with those with latent TB infection (LTBI). MCs respond in vitro to *Mtb* exposure via degranulation and by inducing proinflammatory cytokines. In the current study, we demonstrate an increased production of chymase by MCs in granulomas of humans and macaques with PTB. Single-cell (sc) RNA sequencing analysis revealed distinct MC transcriptional programs between LTBI and PTB, with PTB associated MCs enriched in interferon gamma, oxidative phosphorylation, and MYC signaling. In a mouse model, MC deficiency led to improved control of *Mtb* infection that coincided with reduced accumulation of lung myeloid cells and diminished lung inflammation at chronic stages of infection. Airway transfer of MCs into wild-type *Mtb* infected mice showed increased neutrophils, decreased recruited macrophages, and elevated *Mtb* dissemination to the spleen. Together, these findings highlight MCs as active drivers of TB pathogenesis and potential targets for host-directed therapies for TB.

## Introduction

Tuberculosis (TB) remains a significant global health issue, with approximately one-quarter of the world’s population harboring *Mycobacterium tuberculosis (Mtb)*, causing around 1.25 million deaths each year (WHO, 2024). The disease often starts as a latent TB infection (LTBI), in which the bacteria may remain dormant without disease symptoms. However, LTBI can progress to active pulmonary TB (PTB), characterized by severe respiratory symptoms and high transmission potential. The immune mechanisms that allow progression from latency to PTB are not fully defined. Thus, understanding the immune factors that drive progression toward PTB will allow the development of novel therapeutics for TB control. Towards this overall goal, we recently showed that the lung single-cell transcriptional immune landscape during LTBI and PTB in *Mtb*-infected macaques was distinct. For example, PTB was characterized by the significant accumulation of Type I IFN-expressing plasmacytoid DCs (pDCs), IFN-responsive macrophages, as well as activated T cells in the lungs (Esaulova et al., 2021). Additionally, mast cells (MCs) were increased in the lungs of macaques with PTB (Esaulova et al., 2021). In sharp contrast, LTBI was characterized by increased presence of cytotoxic NK cells but lack of recruitment of MCs in the lungs (Esaulova et al., 2021).

MCs are found in the lung where they influence inflammatory responses (Virk et al., 2016; Wasserman, 1984). MCs have been shown to respond *in vitro* to *Mtb* exposure via surface receptors such as CD48 (Munoz et al., 2003). They also respond to *Mtb* exposure or mycobacterial lipids by undergoing degranulation of prestored granules, such as histamine and P-hexosaminidase, and secrete proinflammatory cytokines such as IL-6 and TNF-a (Munoz et al., 2003). Degranulation of MCs following intratracheal infection with a high dose of *Mtb* was shown to limit inflammation and the production of proinflammatory cytokines such as IL-1β and TNF-α (Carlos et al., 2007). MCs produce and release either chymase or tryptase (Bian et al., 2021), which are both proteases that are stored in the cell’s secretory granules. Recent studies with lung biopsies of TB patients showed an enrichment of MCs expressing IL-17 at inflammatory sites. In contrast, chymase-rich MCs (MCcs) producing TGF-β were detected in proximity to mature granulomas in lung biopsies from PTB (Garcia-Rodriguez et al., 2021). Furthermore, while healthy lung predominantly has tryptase-expressing mast cells (MCTs), both chymase and mast cells co-expressing chymase and tryptase (MCcs and MCTcs) accumulate in the infected lung of patients with PTB (Garcia-Rodriguez et al., 2021). Thus, while previous studies have shown that MCs respond to *Mtb* exposure and accumulate in macaque and human lungs during PTB, it is not completely known if MCs functionally mediate protective or pathological outcomes in the context of TB infection.

In the current study, we showed that the distribution and localization of MCs in PTB in humans and macaques were associated with chymase production. Using scRNA seq analysis, we show that MCs found in LTBI and healthy lungs in macaques are transcriptionally distinct from PTB lungs, showing enrichment of tumor necrosis factor alpha, cholesterol, and transforming growth factor beta signaling. In contrast, MCs found in PTB express increased levels of signatures associated with interferon gamma, oxidative phosphorylation, and MYC signaling. Additionally, mice deficient in MCs showed improved control of *Mtb* infection and reduced lung inflammation. Airway transfer of MCs into wild-type Mtb-infected mice increased lung neutrophils and elevated *Mtb* dissemination to the spleen. These results together provide novel evidence that MCs contribute to immune pathology and reduced *Mtb* control, suggesting a pathological role for MCs during *Mtb* infection.

## Materials and methods

### Study subjects and animal studies

All human lung biopsy samples were obtained from the Tuberculosis Outpatient Clinic and the Department of Pathology at the National Institute of Respiratory Diseases (INER) in Mexico City, before *Mtb* treatment with informed consent, and with the approved protocol by the INER IRB for their use (project numbers B04-15 and B09-23). Also, lung samples from healthy controls (HC), non-TB individuals, were obtained from the tissue repository of the Department of Pathology at INER. No compensation was provided to the patients.

Non-human primate procedures were approved by the Institutional Animal Care and Use Committee of Tulane National Primate Research Center and were performed following National Institutes of Health (NIH) guidelines. Male and female Indian rhesus macaques, verified to be free of *Mtb* infection by tuberculin skin test, were obtained from the Tulane National Primate Research Center. The animals were housed in an ABSL3 facility.

C57BL/6 and B6.Cg-KitW-sh/HNihrJaeBsmJ (Strain #:030764) alias *CgKitW*^*Sh*^ mice were procured from Jackson Laboratory (Bar Harbor, ME) and bred at Washington University in St. Louis or University of Chicago. Six to eight-week-old female and male mice were used in the experiments. All mice were maintained and used per the approved Institutional Animal Care and Use Committee (IACUC) guidelines at Washington University in St. Louis or University of Chicago.

### Aerosol infection

For murine experiments, *Mtb* strain HN878 was cultured in Proskauer Beck medium containing 0.05% Tween 80 until reaching mid-log phase and frozen in 1 ml aliquots at -80°C until used. Mice were aerosol infected with ∼100 colony-forming units (CFU), as described previously (Khader et al., 2007). *Mtb* strain CDC1551 was used to infect NHPs. This species-specific choice reflects differences in pathogenicity: HN878 induces robust disease in mice, while CDC1551, a less virulent strain, allows development of a macaque model that recapitulates latent and chronic TB upon low- to moderate-dose aerosol exposure respectively (Kaushal et al., 2015; Sharan et al., 2021; Singh et al., 2025). This ensures physiologically relevant and controlled studies within each species. Macaques were assigned to three groups: (1) uninfected control, (2) macaques with LTBI were exposed to a low dose (∼10 CFU), and (3) macaques with PTB were exposed to a high dose (∼100 CFU) of *Mtb* CDC1551 via the aerosol route using a custom head-only dynamic inhalation system housed within a class III biological safety cabinet as previously described (Esaulova et al., 2021). The animals were periodically monitored for their physiological parameters and to monitor disease symptoms.

### Bacterial burden and cytokine analysis

Bacterial burden was assessed using serial 10-fold dilutions of lung or spleen homogenates and plated on 7H11 agar solid medium supplemented with OADC (oleic acid, bovine albumin, dextrose, and catalase). Colonies were counted after 2-3 weeks of incubation. Cytokine/chemokine expression was analyzed in lung homogenates from infected mice via Luminex (Millipore-Sigma) or ELISA (R&D) as per the manufacturer’s protocol.

### Generation of single-cell suspensions from tissues and flow cytometry staining

Lung single-cell suspensions from *Mtb-infected* mice were prepared as previously described, (Gopal et al., 2013). Briefly, mice were euthanized with CO2. The right lower lobe was isolated and perfused with heparin in saline. Lungs were minced and incubated with collagenase/DNase for 30 minutes at 37°C. Lung tissue was pushed through a 70µm nylon screen to obtain a single cell suspension. Following lysis of erythrocytes, the cells were washed and resuspended in cDMEM (DMEM high glucose + 10% fetal bovine serum + 1% Penicillin/Streptomycin) for flow cytometry staining. For flow cytometric analysis, cells were either stained immediately or stimulated with phorbol myristate acetate (PMA-50ng/ml; Sigma Aldrich) and ionomycin (750 ng/ml; Sigma Aldrich) in the presence of GolgiStop (BD Pharmingen).

For myeloid cell surface staining, the following fluorochrome-conjugated antibodies were used: CD11b-APC (clone M1/70), CD11c-PE-Cy7 (clone HL3, BD Biosciences), GR-1-PerCP-Cy5.5 (clone RB6-8C5, BD Pharmingen), and MHC class II-FITC (clone M5/114.15.2,Tonb Bioscience), CD117 (cKit)-Super Bright 780 (clone 2BB, eBioscience), and FcεR1-PE (clone MAR-1, eBioscience). Myeloid cell subsets were defined as follows: alveolar macrophages (CDI 1c^+^CDl lb^-^), neutrophils (CDI lb+CDl Ic-Gr-1^hi^), monocytes (CD11b^+^CD11c^−^Gr-1^med^), recruited macrophages (CD11b^+^CD11c^−^Gr-1^low^), and MCs (CDl lb^−^cKit^+^FctεRl^+^). T cells were identified based on a gating strategy as described before (Griffiths et al., 2016)). Surface staining included CD3-AF700 (clone 500A2, BD Biosciences), CD4-Pacific Blue (clone RM4.5, BD Biosciences), CD44-PE-Cy7 (clone 1M7, Tonbo Biosciences), and CD8-APC-Cy7 (clone 53-6.7, BD Biosciences). For intracellular cytokine staining, lung cells were fixed and permeabilized using fixation/permeabilization concentrate and diluent (eBioscience) following the manufacturer’s instructions. Cells were then stained with IFNγ -APC (clone XMG1.2, Tonbo Biosciences) and TNF-α-FITC (clone MP6-XT22, BD Pharmingen), or respective isotype controls (APC rat IgG1κ and FITC rat IgG1α, BD Pharmingen) for 30 minutes. Samples were acquired on a four-laser BD Fortessa Flow Cytometer, and data were analyzed using FlowJo software (Treestar). Absolute cell numbers for each population were back calculated based on total viable cell counts per lung sample.

### In vitro culture and intratracheal delivery of MCs

Six to eight weeks old C57BL/6 mice were euthanized by CO_2_, and femurs and tibia were collected. Bone marrow cells obtained after RBC lysis were suspended in Bone Marrow Derived MC (BMMC) media containing RBMI media (Gibco) supplemented with 20% FBS (Sigma), 4mM-glutamine (Sigma), 25mM HEPES (Corning), 5 × 10^−5^ 2-mercaptoethanol (Sigma), lmM sodium pyruvate (Sigma), 0.lmM nonessential amino acids (Gibco), Penicillin and Streptomycin (Sigma), 20ng/ml murine IL-3 and 20ng/ml murine Stem Cell factor (Peprotech) and maintained in BMMC media at 1 × 10^6^ cells per ml in T75 flasks at 37°C in 5% CO2 incubator. Cells were fed twice every week with fresh BMMC media by centrifuging non-adherent cells at 1000 rpm for 10 min at room temperature, resuspending at a density of l × l0^6^/ml, and were maintained for 30 days, more than 95% cells were positive for FcεRI and c-kit, which was confirmed by flow cytometry (Varma & Puri, 2019). 5 × l0^4^ BMMC having 95% FcεRI and c-kit positivity were transferred in C57B1/6 through the intratracheal route.

### Morphometric analysis of lung Histopathology and neutrophil inf ltration

For mouse studies, the left upper lobe was collected for histomorphometric analysis. The lobes were infused with 10% neutral buffered formalin and embedded in paraffin. 5 µm-thick lung sections were cut using a microtome, stained with hematoxylin and eosin (H&E), and processed for light microscopy. Images were captured using the automated Nanozoomer digital whole slide imaging system (Hamamatsu Photonics). Regions of inflammatory cell infiltration were delineated utilizing the NDP view2 software (Hamamatsu Photonics), and the percentage of inflammation was calculated by dividing the inflammatory area by the total area of individual lung lobes. All scoring was conducted in a blinded manner. Formalin-Fixed Paraffin Embedded (FFPE) lung sections were also stained with APC-conjugated rat anti-mouse Ly6G (clone 1A8, BioLegend, RRID:AB_2227348). Nuclei were counterstained with DAPI. Neutrophils were quantified in three randomly selected 200× fields per lung section. Images at 200× magnification were acquired using a Zeiss Axioplan microscope and recorded with a Hamamatsu camera.

FFPE lung sections from healthy individuals, tuberculosis patients, and NHP infected with *Mtb* were stained with goat anti-human mast cell chymase (LifeSpan Biosciences, LS-B4134, RRID: AB_l0718418) and rabbit anti-human tryptase (Cell Signaling Technology, 195235). Primary antibodies were detected with Alexa Fluor 568 donkey anti-goat IgG (Thermo Fisher Scientific, A-11057, RRID: AB_2534104) and Alexa Fluor 488 donkey anti-rabbit IgG (Jackson ImmunoResearch Laboratories, 711-546-152, RRID: AB_2340619). Nuclei were labeled with DAPI. MC positive for chymase, tryptase, or both were blindly quantified in three 200× random fields per sample in human and NHP lung sections. 200× pictures were taken with an Axioplan Zeiss microscope and recorded with a Hamamatsu camera.

### Single-cell data reanalysis

The NHP single cell lung data was accessed from GEO (GSE149758) and processed through Cell Ranger v7.0 using the Macaca Mulatta reference genome (Mmul_l0). The obtained matrix file was processed through the R package *Seurat v5* for downstream analysis of the count matrix. The cells were filtered based on mitochondrial gene content and were selected for analysis when at least 500 genes were detected. Data was log normalized. The most variable genes were detected by the *FindVariableFeatures* function and used for subsequent analysis. Latent variables (number of UMIs and mitochondrial content) were regressed out using a negative binomial model (function *ScaleData)*. Principal component analysis (PCA) was performed with the *RunPCA* function. A UMAP dimensionality reduction was performed on the scaled matrix (with most variable genes only) using the first 20 PCA components to obtain a two-dimensional representation of the cell states. For clustering, we used the functions *FindNeighbors* (20 PCA) and *FindClusters* (resolution 0.25). MCs were identified and re clustered based on expression of the canonical MC marker genes *FCER1A* (High affinity Fe IgE receptor), *CD48, FCER1G* (Fe IgE receptor), *MS4A2* (IgE subunit), and *ITGAX* (CDI le) as a negative marker. The cells identified MC cluster (only one cluster) were subset and re clustered using the method outlined above at a resolution of 0.1. To identify marker genes for mast cells, we used *FindAllMarkers* to compare the cluster against all other clusters and *FindMarkers* to compare selected clusters. For each cluster, only genes that were expressed in more than 15% of cells with at least 0.15-fold differences were considered. The differential genes were subjected to enrichment analysis using the Hallmark Pathway gene set from *MsigDB*. Only the pathways that met an FDR threshold less than 0.05 were considered. Gene signatures were defined with the R package *Ucell*. The output is a module signature score generated by the *AddModuleScore* function. The obtained score was overlaid on the UMAP and visualized. The Values per cell were extracted and used to plot a summed module U cell score. GraphPad Prism was used for the Violin plots and the heatmap. All other figures were generated in R.

An independent *M.fascicularis* lung single-cell RNA-seq dataset (GSE200151) (Gideon et al., 2022) was downloaded and processed using Cell Ranger v7.0 with the *Macaca mulatta* reference genome (Mmul_l0). The resulting count matrices were imported into Seurat v5 for downstream analysis. Cells were filtered based on mitochondrial gene content and were retained if they expressed at least 500 genes. Data were log-normalized, the most variable genes were identified with *FindVariableFeatures*, and confounding effects of sequencing depth (UMI counts) and mitochondrial fraction were regressed out during scaling with ScaleData. Principal component analysis (PCA) was performed, followed by clustering *(FindNeighbors, FindClusters)* and UMAP embedding *(RunUMAP)*. MCs were identified and subset based on canonical marker expression *(FCER1A, CD48, FCER1G*, *MS4A2)* and re-clustered at low resolution to ensure purity of the MC population. To quantify MC subsets across disease severity states, we culated the proportion of cells expressing chymase (*CMA1*), tryptase *(LOC102140229, TPSG1)*, or dual-positive *CMA1*^*+*^*LQC102140229*^*+*^ using binary thresholds (>0 counts). Fisher’s exact tests were performed to compare proportions between MCs from high-burden granulomas (4 weeks, more severe disease; n = 372 MCs) and low-burden granulomas (10 weeks, less severe disease; n = 7,306 MCs). For each comparison, we reported the odds ratio (OR) with 95% confidence interval and the exact p-value. To examine functional programs, we computed UCell signature scores for Hallmark IFNγ; signaling, TNF signaling, and oxidative phosphorylation. Signature score distributions were compared between severity groups. All statistical tests were performed in R (v4.3.l) with the Seurat (v5.0) and UCell (v2.0) packages. Violin and bar plots were generated in R or exported to GraphPad Prism for visualization.

### Data analysis and statistics

All data were analyzed using the indicated methodology in each figure legend. Two-sided unpaired t-test was performed for comparing the significance between 2 groups, one-way ANOVA, Tukey’s test, and two-way ANOVA Sidak’s multiple comparison test was performed for more than 2 groups using GraphPad Prism 5 and 10, respectively (La Jolla, CA). Significance is denoted on the figure and the respective figure legends. Outliers, if any, were removed using Grubb’s outlier test and mentioned in the respective figures.

## Results

### Mast cells localize and transtion phenotypes within TB granulomas

A previous study observed that tryptase-expressing MCTS were primarily found within the lungs of healthy controls (HC), while chymase-expressing MCcs or both chymase and tryptase-expressing MCTcs were found in the lung of patients with PTB (Garcia-Rodriguez et al., 2021). To corroborate these observations and analyze the compartmentalization of MCs in human lungs, we stained lung biopsies from healthy individuals and patients with PTB to visualize the spatial distribution of MC_T_s, MC_C_s, and MC_TC_S. Lung granulomas from PTB patients were further classified by the presence or absence of necrosis, as early or late granulomas. Early granulomas had well-defined immune and stromal cell clusters without central necrosis. Late granulomas were larger, with necrotic cores containing bacteria and dead neutrophils, surrounded by lymphocytes. MCs were quantified both within and around early granulomas, whereas in late granulomas, they were primarily measured in the peripheral regions surrounding the necrotic center.

Considering lung parenchyma, interstitium, vasculature, or bronchus, we observed that HC lungs predominantly contain MC_T_s and, to a lesser extent, MC_TC_s. In TB lesions from PTB patients, MC_TC_S accumulated in early immature granulomas, whereas MC_C_s accumulated in late granulomas (Figure 1A-B). MC_T_S also increased in the interstitium, vasculature, and bronchus-associated lymphoid tissue of PTB patients (Figure S1A-C). However, we did not observe any differences in MC_TC_s and MC_C_s at these sites (Figure S1D-I). These observations confirmed that tryptase-expressing MC_T_S are found in HCs (Garcia-Rodriguez et al., 2021), while the dual tryptase and chymase-expressing MCs were seen in early granulomas, and only chymase-associated MCs were observed in late granulomas with necrotic cores, defined the MCs associated with TB disease progression.

**Figure 1.**
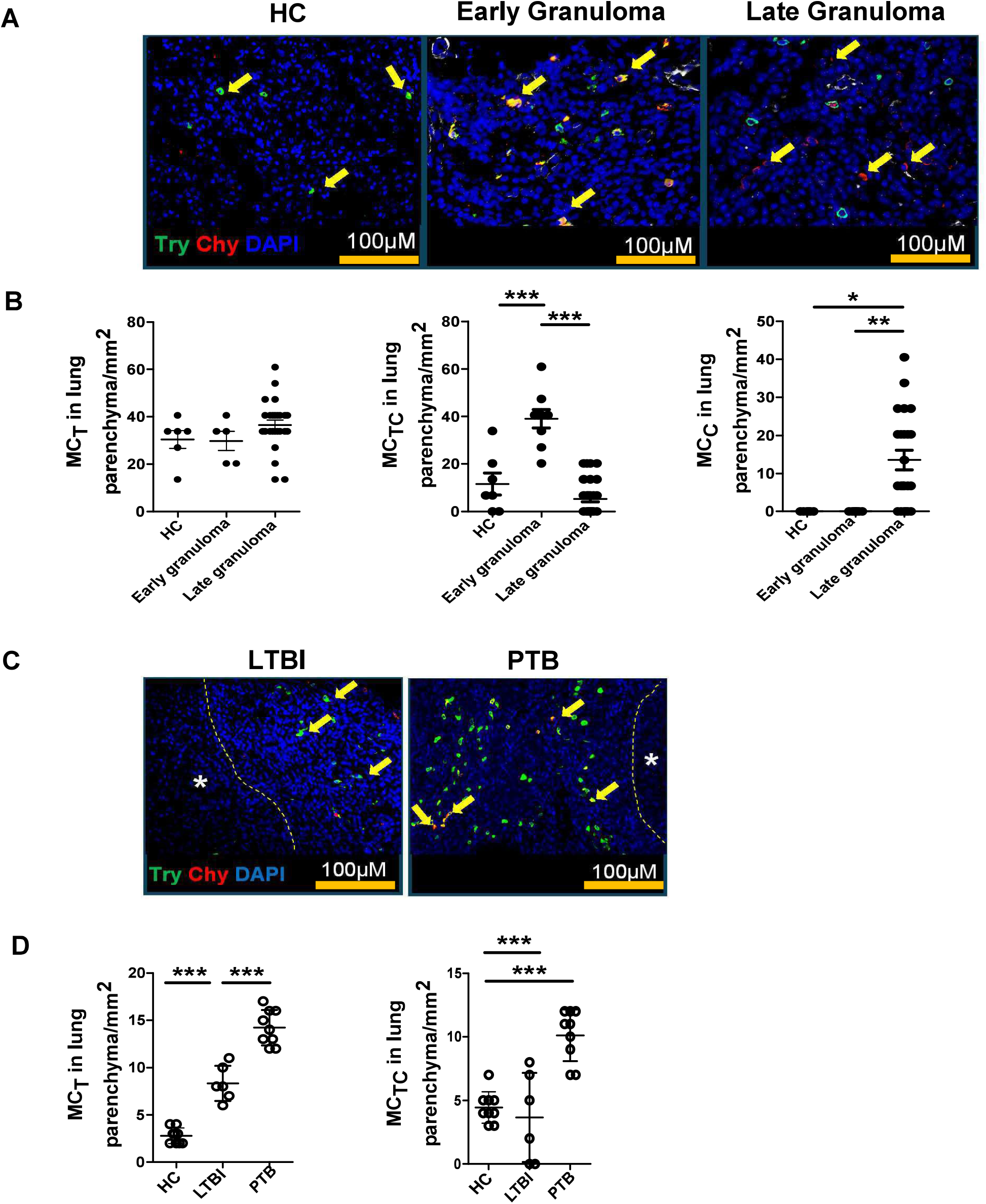
Chymase positive MCs are predominant in TB infected human and macaque lung tissue. Lung biopsies from healthy individuals (n = 4) or patients with PTB (n = 5) were stained for tryptase MC_T_ (green) or chymase MC_c_ (red). (A) Immunofluorescence microscopy shows MC_TS_ (green) in healthy lung biopsies (HC). MC_Tcs_ (red and green merge) are located around the early granulomas, while MC_CS_ (red) surround the late granulomas in TB infected lung biopsies. (B) Predominance of MC_TS_ in healthy lungs transitioning to MC_TCS_ in early granuloma and becoming MC_CS_ in late granulomas in TB infected lungs. (C) Immunofluorescence microscopy shows MC_TS_ (green) and MC_TCS_ (merge) in lungs of healthy (HC), LTBI and PTB macaques. (D) Predominance of MC_TS_ (green) and MC_TCS_ (merge) in PTB compared to LTBI and HC. Statistical analysis was performed using unpaired, 2-tailed Student’s t test,**** p < 0.0001, *** p < 0.001, * p< 0.05.

Our previously published data showed that MCs accumulate in the lungs of macaques with PTB compared to LTBI (Esaulova et al., 2021). Thus, we next analyzed the accumulation and localization of MCs in the lungs of macaques with LTBI and PTB. We found that, similar to human healthy lungs, MC_T_S accumulated in the lungs of healthy macaques. Although MC_T_S increased in some lesions in the lungs of macaques with LTBI, the numbers of MC_T_s in macaques with PTB were significantly increased in all sites, including the granuloma (Figure 1C-D), interstitium, vasculature, as well as bronchus-associated lymphoid tissue of PTB patients (Figure S1J-L). Additionally, MC_TC_S were significantly increased within the granulomas of macaques with PTB compared to the lungs of macaques with LTBI and HCs (Figure 1C-D), but did not differ at any other sites in the lung (Figure S1M-O) compared to HCs. However, we did not observe any increase in these cells at other sites within the lung compared to healthy macaques. MC_C_s were not measurable in any region of the macaque lungs. Our data indicate that during LTBI, there is an accumulation of MC_T_s but not MC_TC_s. However, as the disease progresses to PTB, both MC_TC_s and additional MC_T_s are elevated, particularly within the granulomatous lesions.

### Lung single-cell transcriptome in macaques with tuberculosis exhibits MC diversity

In the previous section, we found that tryptase protein expression on MCs was lower in LTBI, and as the disease progressed to PTB, MCs expressed chymase, either alone MC_C_s or in combination with tryptase, MC_TC_. To further validate whether this increase in tryptase and chymase protein expression was also reflected in the single-cell transcriptomes during PTB, LTBI, and HCs, we re-analyzed the MCs from our previously published data from NHPs (Esaulova et al., 2021). These macaques were infected with 10 bacilli to generate LTBI, or with 100 bacilli for a progressive PTB infection, or were uninfected (Figure 2A). The median duration of infection for the PTB macaques was 10 weeks, and the LTBI macaques underwent necropsy at a median of 23 weeks post-infection. We analyzed the single-cell transcriptomes of 500 MCs using unsupervised clustering and identified four distinct clusters. Three clusters (0, 1, and 3) belonged to the PTB group, while cluster 2 was found exclusively in LTBI and HC, with the majority of MCs coming from the PTB condition (Figure 2B). All the MC clusters were positive for canonical markers such as *FCER1A* (high-affinity lgE receptor), *MS4A2* (lgE subunit), *CD48* (mast cell receptor), and negative for markers like *ITGAX* (macrophage/dendritic cell marker) (Figure S2A), with distinct differentially expressed genes (DEGs) (Figure S2B). Reactome gene ontology analysis of cluster-specific DEGs revealed enrichment of cholesterol, TNF-α, and TGFβ signaling in LTBI, while oxidative phosphorylation, IFNγ signaling, and MYC signaling were enriched in the PTB group (Figure 2C). Plotting the summed *Ucell* module scores revealed significant upregulation of IFNγ signaling, oxidative Phosphorylation, and Th2 signature in PTB (P < 0.05), while LTBI and HC clusters showed enhanced TNF-α signaling (P <0.05) (Figure 2D-G; Figure S2 C-F). These results indicate that MCs exhibited distinct transcriptomic profiles depending on the disease state, with MCs from LTBI and HC showing more metabolically active pathways, while MCs from PTB display a more pro-inflammatory signature.

**Figure 2.**
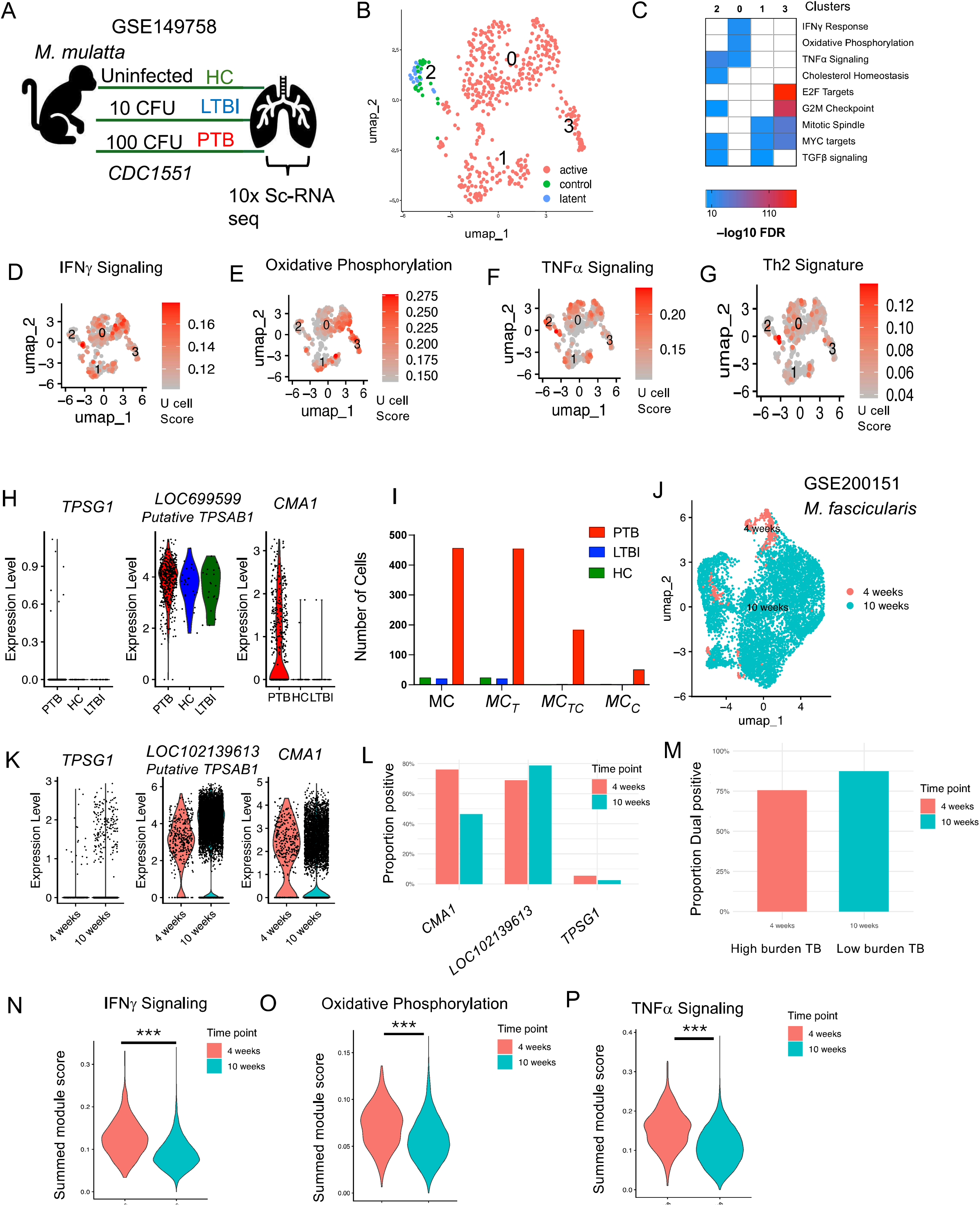
MC signatures across disease conditions in NHPs. Data was re-analyzed from the lungs of *M. mulatta* infected *withMtb CDCJ551* (GSE200151). (A) Schematic of the study design across disease conditions (B) UMAP embedding of *FCER1A+* mast cells, showing the distribution of these cells across the different disease conditions (PTB in pink, HC in green, and LTBI in blue). (C) Heatmap of Hallmark pathway analysis for differentially expressed genes, highlighting the top pathways with the highest FDR values for each condition. (D-G) UCell module for pathways: IFNy signaling (D), TNF-*α* signaling (E), Oxidative Phosphorylation (F), and Th2 signature (G) across disease conditions, shown on UMAP embeddings. (H) Violin plots of gene expression for key MC markers *(CMA1, TPSG1, LOC699599)* across disease conditions. (I) Cell counts of different MC subtypes (MC_C_, MC_T_, MC_TC_) across disease conditions (PTB, red bars, LTBI, blue bars, and HC green bars). (J) UMAP plot of the NHP lung granuloma dataset (GSE200151), showing the distribution of cells at 4 weeks (high disease burden) and 10 weeks (low disease burden) in *M*.*fasicularis* infected with *Mtb Erdman*. (K) Gene expression violin plots for key MC markers *(CMA1, TPSG1, LOC699599)* from the new dataset across time points. (L) Proportions of different MC subtypes (MC_C_, MC_T_, MC_TC_). (M) Violin plots of summed module scores for the key pathways (IFN*γ* signaling, TNF-*α* signaling, oxidative phosphorylation) across disease burdens, showing pathway activity. Statistical significance was assessed using Kruskal-Wallis tests with Dunn’s multiple comparison correction (**p < 0.01, ***p < 0.001, ****p < 0.0001).

Since we observed increased tryptase and chymase in MCs of PTB macaques (Figure 1C), we next examined the levels of tryptase and chymase genes within the single-cell dataset. As we were examining these genes across species, we observed considerable variation in sequence similarity and functional annotation of tryptase genes between humans and NHPs. While *TPSG1* (encoding γ tryptase) and *TPSD1* (encoding δ tryptase) share the gene name in humans and NHPs, the gene corresponding to the more widely expressed *TPSAB1* (encoding α and β1 tryptase) differs in NHPs. Based on phylogenetic similarity to human α and β tryptase, the NHP ortholog is often referred to as α-like or β-like (α/β) tryptase. However, their gene names differ, as they are still classified as putative proteins and not formally annotated with the same nomenclature as in humans. The putative tryptase genes in NHPs are annotated as *LOC699599* for *Macaca (M). mulatta* and *LOC102139613* for *M.fasicularis*. Examining these genes in the NHP single cell transcriptome dataset, we detected the expression of the γ and the putative α/βtryptase genes, but found no expression for δ tryptase. *TPSG1* was found to be expressed at low levels and only in a few MCs from the PTB group. In contrast, the putative α/β tryptase gene was expressed in MCs across all groups, with the highest expression in PTB (Figure 2H), consistent with our immunofluorescence data (Figure 1D). As expected, the chymase (encoded by *CMA1)* expressing cells were detected exclusively in the PTB group and were absent in LTBI and HCs (Figure 2H). To confirm whether these *CMA1*-positive cells also co-expressed the putative α/β tryptase as well, we quantified cells expressing single and dual transcripts. This analysis revealed that most of the chymase-positive cells (243 cells) also expressed the putative α/β tryptase gene (183 cells), supporting our earlier observation of a dual tryptase chymase MC signature linked with PTB. In contrast, MCs from HC and LTBI groups showed expression of tryptase alone (Figure 2I).

To strengthen our findings, we validated MCs using an independent lung single-cell transcriptome from NHP *(M.fasicularis)*, collected at 4 weeks (higher bacterial load, more severe disease) and 10 weeks (low bacterial load, less severe disease) following low-dose *Mtb* Erdman infection (Gideon et al., 2022; Grimbaldeston et al., 2005). This dataset had 372 MCs at 4 weeks, and 7306 MCs at 10 weeks (Figure 2J). We examined the expression of chymase and several tryptase genes, including *TPSG1, TPSD1*, and *LOC102140229* (putative NHP ortholog for human *TPSAB1)*. While *TPSD1* expression was undetectable, the other tryptase genes showed high expression with a pronounced increase in chymase *(CMA1)* expression both at 4 and 10 weeks (Figure 2K). Similar to our transcriptomic scRNA seq dataset, we quantified MCs co-expressing *LOC102140229* and *CMA1*. Consistent with our data, *CMA1* and *LOC102140229* (the α/β -like tryptase ortholog) expressions were associated with severe disease, being significantly enriched in MCs from the higher-burden 4-week granulomas, (OR = 0.27, *p* < 1 × 10^−29^, and OR= 1.68, *p* < 1 × 10^−5^, respectively). Importantly, when restricting analysis to *CMA1*^*+*^ cells, the dual-positive *CMA1*^*+*^*LOC102140229*^*+*^ MC subset was proportionally more abundant in less severe, 10-week granulomas (OR = 0.51, *p* < 1 × 10^−9)^ (Figure 2M). This suggested that while chymase *(CMA1)* expression marked severe disease, the presence of the dual tryptas—chymase phenotype in less severe lesions, supported the idea that MC transcriptional diversity emerges in association with disease modulation This further supporting our observation of MC diversity with increasing dual tryptase and chymase signature associated with disease progression. Similar to our observations in *M.mulatta*, we quantified the DEGs and carried out Ucell scores in the MC subset from *M.fascicularis*.We observed that MC from high burden TB granulomas showed higher IFNγ signaling, oxidative phosphorylation (Figure 2N-O), similar to higher PTB scores seen in *M.mulatta*. However, unlike *M.mulatta, M.fascicularis* also showed increased TNF signaling in high-burden granulomas (Figure 2P). These results highlight that MCs in high-burden granulomas upregulated IFNγ, TNF, and metabolic programs, coupled with chymase expression, whereas less severe granulomas were enriched for tryptase-positive MCs. Although this mirrors our findings seen in in *M.mulatta*, we observed species-specific differences in how TNF signaling is distributed across disease states.

### MC-deficient mice exhibit enhanced control of *Mtb*

We next determined whether MCs are induced in response to *Mtb* infection in mice, and characterized their accumulation early and later in infection. In our previous studies, we observed that innate cells such as innate lymphoid cells (ILCs) accumulate rapidly between days 5-10 post-infection, followed by neutrophils, macrophages, and monocytes between days 10-15, and T cells by days 21 to 30 (Ardain et al., 2019). We found that MCs begin to accumulate in the lungs between days 21 and 30, coinciding with the timing of *Mtb* growth (Figure S3A-B) and T cell recruitment. To investigate the functional role of MCs in *Mtb* infection, we utilized the MC-deficient mouse model, *CgKitW*^*Sh*^. These mice carry spontaneous loss-of-function mutations in both alleles of the dominant *white spotting* (W) locus *(i*.*e*., *c-kit)*, leading to impaired *c-kit* tyrosine kinase-dependent signaling resulting in dysregulated MC development, survival, and function (Wolters et al., 2005). We infected *CgKitW*^*Sh*^ mice with low-dose aerosolized *Mtb* strain HN878 for early (50 days post infection, dpi) and later time points (100 dpi and 150 dpi) and compared them with C57BL/6 wild-type (WT) Mtb-infected mice (Figure 3A). While there were no significant differences observed in lung and spleen bacterial burden at 50 dpi between *CgKitW*^*Sh*^ and WT *Mtb*-infected mice, *CgKitW*^*Sh*^ *Mtb* infected mice showed significantly lower lung and spleen *Mtb* CFU compared to WT *Mtb* infected controls at 100 dpi. However, at 150 dpi, lung bacterial burden in *CgKitW*^*Sh*^ *Mtb* infected mice trended lower but not significant, these mice showed enhanced bacterial control in the spleen (Figure 3B-C). The reduction in bacterial load also coincided with reduced lung inflammation in *CgKitW*^*Sh*^ *Mtb-infected* mice at 150 dpi (Figure 3D-E). These findings indicate that MC-deficient *CgKitW*^*Sh*^ mice exhibit improved control of *Mtb* infection during chronic stages, suggesting that MCs may contribute to disease persistence or pathogenesis in chronic TB.

**Figure 3:**
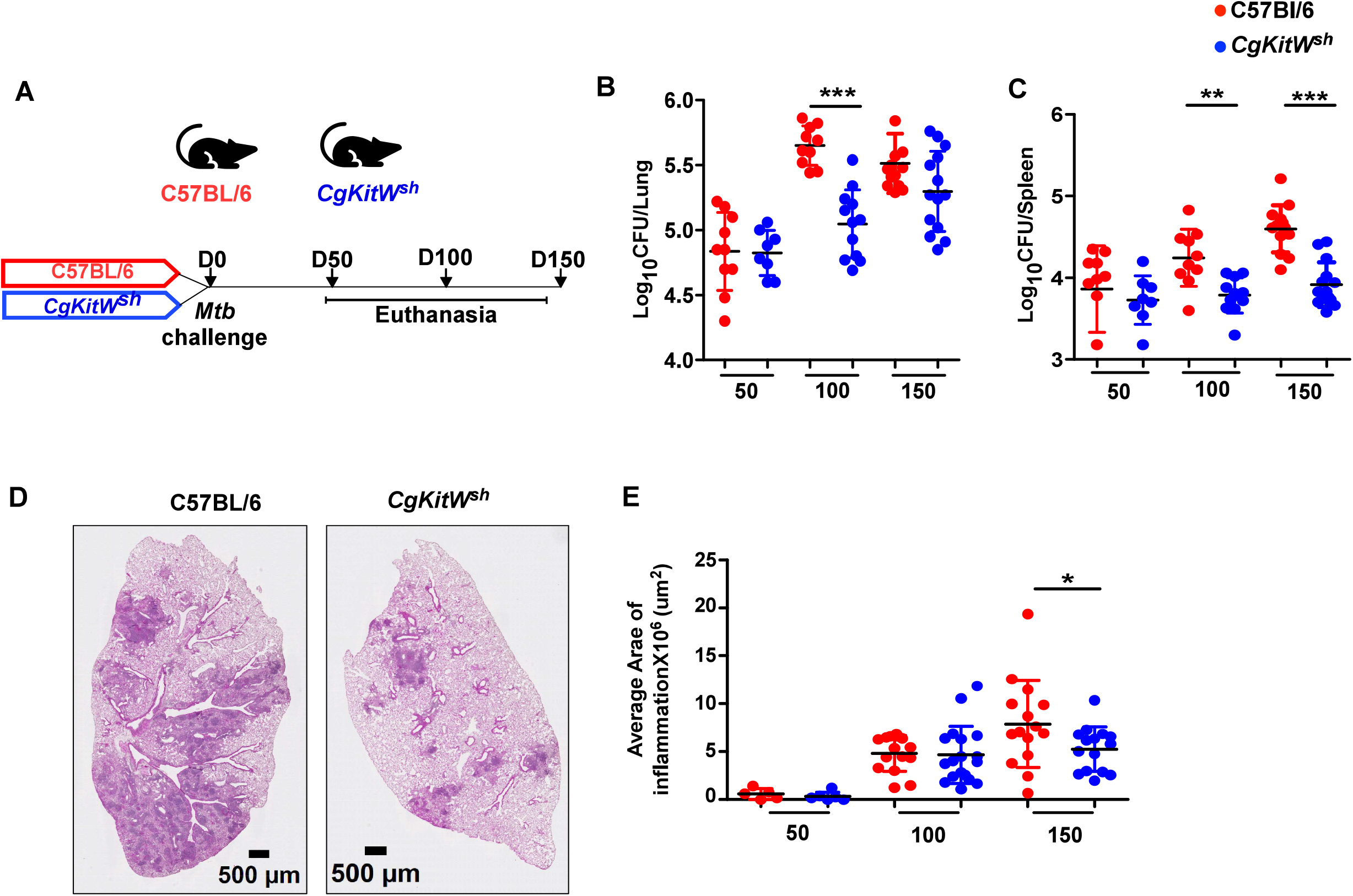
MC-deficient mice are resistant to *Mtb* chronic infection. (A) C57BL/6 and CgKitW^sh^ mice were infected with a low aerosol dose (∼l00CFU) of *Mtb* HN878 and mice were sacrificed at 50, 100 and 150 dpi. (B) Bacterial burden was assessed in lungs and spleens by plating. (C) Lungs were harvested, fixed in formalin and embedded in paraffin. H&E staining was carried out for blinded and unbiased analysis of histopathology. (D) Representative images and the area of inflammation measured in each lobe are shown. Scale bars: 2mm. Original magnification: ×20. Data points represent the mean ± SD of two experiments *(n* = 8-15 per time point per group). Statistical analysis was performed using unpaired, 2-tailed Student’s t test between C57BL/6 and *CgKitW*^*Sh*^ mice, **** p < 0.0001, *** p < 0.001, * p< 0.05.

### MC-deficient mice display altered innate immune cell profiles during chronic *Mtb* infection

To further address the functional basis of enhanced protection observed in MC-deficient mice, we analyzed the lung immune responses in *CgKitW*^*Sh*^ mice both at baselines and following *Mtb* infection, given that this mouse strain is associated with other known immune alterations (Grimbaldeston et al., 2005). MCs were significantly reduced in the lungs of *CgKitW*^*Sh*^ mice compared to WT mice at baseline (Figure 4A). However, we did not observe any significant differences in other innate immune populations in the lung of *CgKitW*^*Sh*^ mice, including dendritic cells (DCs), recruited macrophages (RMs), alveolar macrophages (AMs), neutrophils, monocytes, and T cells at baseline compared to WT controls (Figure 4B-F and S4). Following *Mtb* infection, MCs accumulated progressively in the lungs of both WT and *CgKitW*^*Sh*^ mice up to 100 dpi, after which their numbers stabilized through 150 dpi. In contrast, MC accumulation was significantly impaired in *CgKitW*^*Sh*^ mice throughout infection (Figure 4A). At 50 dpi, RMs were elevated in *CgKitW*^*Sh*^ mice, with no significant difference observed in DCs or neutrophils compared to WT mice (Figure 4B). By 100 dpi, both DCs and neutrophils were decreased in *CgKitW*^*Sh*^ mice; however, these changes were not sustained at 150 dpi (Figure 4C-D). Across all time points, AMs and monocyte populations remained comparable between *CgKitW*^*Sh*^ and WT *Mtb-infected* mice (Figure 4E-F). Previous studies have implicated MCs in driving T cell responses (Elieh Ali Komi & Grauwet, 2018), therefore, we next examined T cell responses induced post-infection. We found no differences in the activated CD4^+^ and CD8^+^T cell responses at 50 dpi however, by 100 dpi, both populations were significantly reduced at 100 dpi in *CgKitW^wh^ Mtb*-infected mice (Figure S5A and E). This reduction extended to functional subsets, with fewer CD4^+^T cells producing IFNγ, as well as diminished dual TNF-α and IFNγ producing cells in the *CgKitW*^*Sh*^ *Mtb-infected* mice as compared to WT *Mtb-infected* mice (Figure S5B-D). We did not find any significant differences in the CDS^+^T cells producing IFNγ, TNF-α, and dual IFNγ and TNF-α producing cells in the *CgKitW*^*Sh*^ *Mtb-infected* mice (Figure S5F-H). Finally, we measured cytokine responses in the lungs of *CgKitW*^*Sh*^ and WT *Mtb-infected* mice at 150 dpi. We found that pro inflammatory cytokines that direct monocyte/macrophage and T cell responses (G-CSF, IFNγ, IL-1, IL-6, IL-17, MCP-1, TNF-α, and RANTES) were significantly lower in *CgKitW*^*Sh*^ *Mtb* infected lungs compared to WT *Mtb-infected* lungs. Similarly, chemokines driving neutrophil recruitment (MIP-1β, and MIP-2) were reduced in *CgKitW*^*Sh*^ *Mtb-infected* mice, whereas Th2 cytokines such as IL-13 did not differ between the two groups (Figure 4G). Together, these results provide evidence that MCs are induced following *Mtb* infection, accumulate in the lung, and mediate immune responses that drive pathology and promote *Mtb* susceptibility and dissemination during chronic TB.

**Figure 4:**
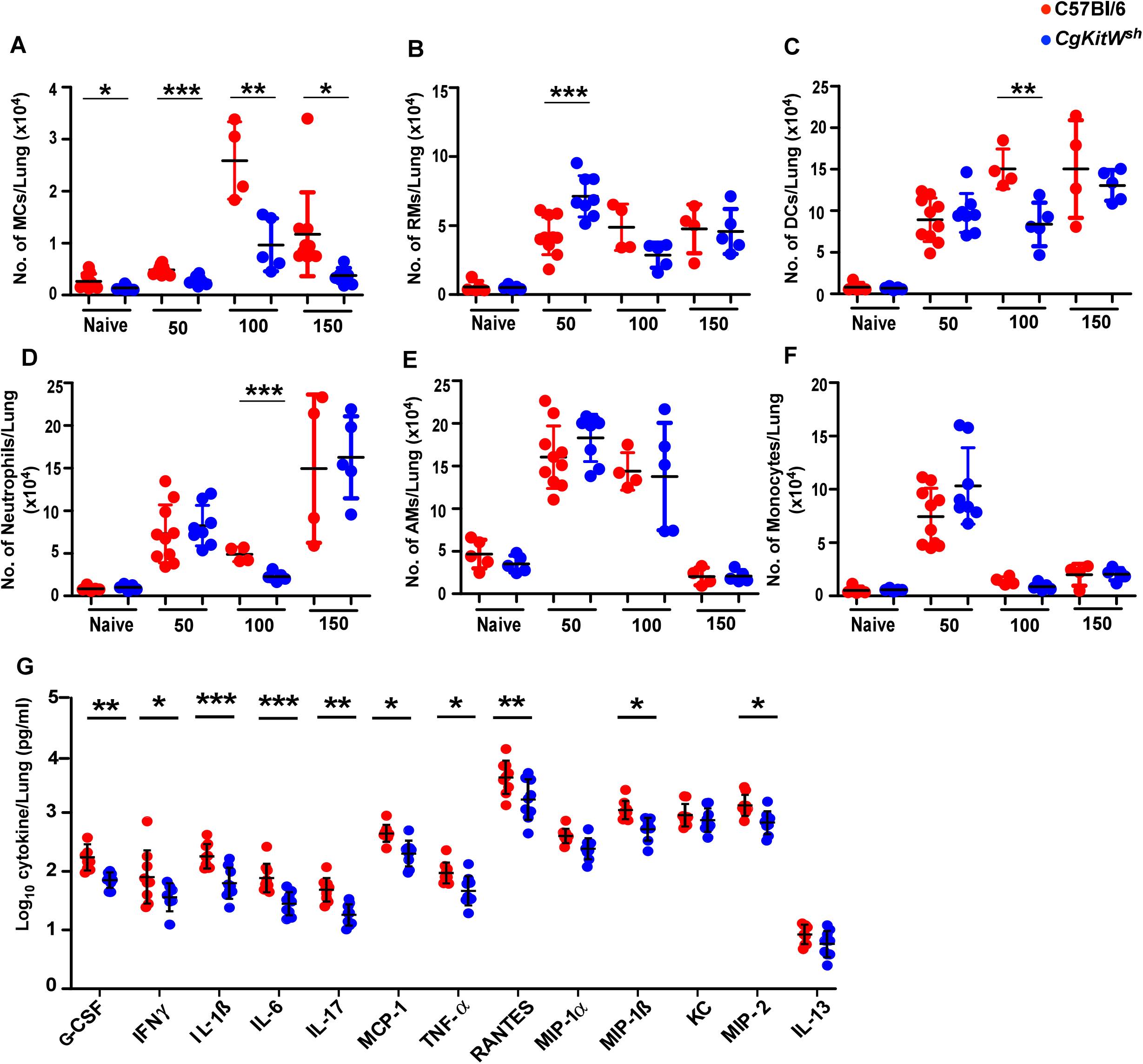
MC-deficient mice have dysregulated immune profile after *Mtb* infection. C57BL/6 and *CgKitW*^*Sh*^ mice were infected with a low aerosol dose (∼l00CFU) of *Mtb* HN878 and mice were sacrificed at 50, 100 and 150 dpi. Number of (A) MCs, (B) DCs (C) RMs, (D) neutrophils (E)AMs, and (F) monocytes were enumerated in the lungs of *Mtb-infected* mice. (G) Cytokine and chemokine production in lung homogenates from mice, collected at 150 dpi, was assessed by multiplex cytokine analysis. Data points represent the mean ± SD of 1 of 2 individual experiments *(n* = 4-10 per time point per group). Statistical analysis was performed using unpaired, 2-tailed Student’s t test for (A) to (F) and Two-way ANOVA Sidak’s multiple comparison test for (G) between C57BL/6 and *CgKitW*^*sh*^ mice, *** p < 0.0001, ** p < 0.001, * p< 0.05. Outliers were removed from the subsets using Grubb’s outlier test.

### Transfer of MCs into lung airways promotes neutrophil accumulation and *Mtb* dissemination

To examine whether MCs can impact *Mtb* control and dissemination, we adoptively transferred bone marrow-derived MCs into the airways of WT mice, hereafter referred to as B6^MC^ mice. A total of 5× l0^4^ MCs were transferred, approximating the number of MCs present in the lungs of WT *Mtb-infected* mice at 100 dpi. Following transfer, mice were infected with *Mtb*, and we found that MCs were retained in the lungs of B6^MC^ *Mtb-infected* mice up to 30 dpi (Figure 5A). Strikingly, B6^MC^ *Mtb-infected* mice showed increased neutrophils frequencies and reduced RMs when compared with B6 *Mtb-infected* mice (Figure 5B, and C). However, no differences in the total numbers of lung MCs, neutrophils, RMs or DCs were observed in B6 and B6^MC^ groups (Figure S6). While the presence of additional MCs in the lungs did not alter the bacterial burden *(Mtb* CFU) and inflammation in the lungs, B6^MC^ *Mtb-infected* mice showed a significant increase in the *Mtb* CFUs in the spleen, suggesting a role for MCs in promoting dissemination from the lungs (Figure 5D-F). Consistently, histological analysis revealed greater neutrophil infiltration in lung sections of B6^MC^ *Mtb-infected* mice compared to WT *Mtb-infected* mice (Figure 5G and H), implicating MCs in driving neutrophil recruitment and potentially enhancing the systemic spread of *Mtb*.

**Figure 5:**
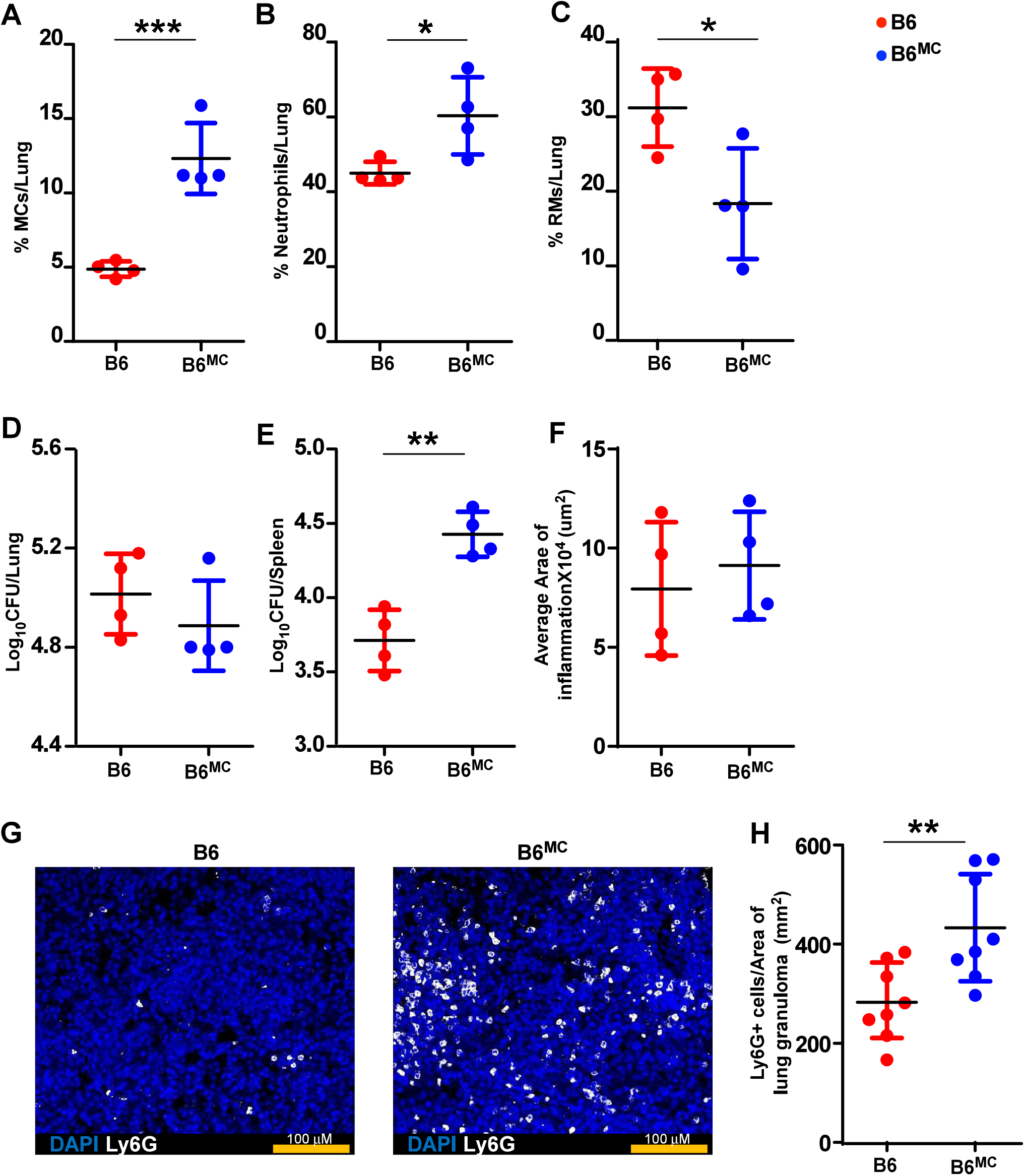
Wild-type mice with airway transferred MCs promote bacterial dissemination. Bone marrow derived in vitro cultured MCs (5×l0^4^ cells/mouse) were adoptively transferred into the lung airways of C57BL/6 mice 7 days before infecting with a low aerosol dose (∼l00CFU) of *Mtb* HN878. MCs were replenished in these mice at 15 dpi, and mice were sacrificed at 30 dpi. Frequencies of (A) MCs, (B) neutrophils, and (C) RMs were enumerated in the lungs of *Mtb-infected* mice. Bacterial burden was assessed in (D) lungs and (E)spleens by plating. (F) Lungs were harvested, fixed in formalin, and embedded in paraffin. H&E staining was carried out for blinded and unbiased analysis of histopathology. (G) Immunofluorescence microscopy shows more neutrophil infiltration in the lungs of MC transferred WT mice. (H) Ly6G^+^ cells per area of lung granuloma measured in each lobe are shown. Scale bars: 2mm. Original magnification: ×20. Data points represent the mean± SD. Statistical analysis was performed using an unpaired, 2-tailed Student’s t test between the groups, **** p < 0.0001, *** p < 0.001, * p< 0.05.

## Discussion

The immune mechanism(s) that mediate the progression from LTBI to PTB are unclear. In this study, we identified MCs as an innate cell type that is overrepresented during PTB, transcriptionally expressed signatures associated with IFNγ, oxidative phosphorylation, and MYC signaling, and localized within mature TB granulomas. Importantly, using mice deficient in MCs, we showed a potential pathological role for MCs in mediating susceptibility to TB, thus providing MCs as a novel therapeutic target.

MCs have been shown to interact with *Mtb* through the GPI-anchored molecule CD48 (Munoz et al., 2003), interaction with TLR2 (Carlos et al., 2007), and potentially TLR4 (Mccurdy et al., 2001). Additionally, *Mtb* is also thought to be internalized by lipid rafts on MCs (Munoz et al., 2009), thus serving as a long-lasting reservoir for *Mtb* (da Silva et al., 2014). These in vitro studies have shown that MC exposure to *Mtb* results in degranulation of MCs, as well as the induction of proinflammatory cytokines such as TNF-α and IL-Iβ. Consistently, another study using the same route of high-dose infection reported increased bacterial burden in both the lungs and spleen of MC-deficient Kit^W-sh/W-sh mice compared to wild-type C57BL/6 controls (Villareal-Rivota et al., 2025). While these studies using a high-dose model of infection reported early induction of inflammatory mediators from MC within hours to days, our in vivo results using a physiological low dose of *Mtb* infection model showed that MCs accumulated between 21-30 days, coinciding with the onset of T cells in the lung. Interestingly, despite the accumulation of MCs in the lung at 30 dpi following low-dose aerosol infection, the impact of MC deficiency on *Mtb* control and inflammation in *CgKitW*^*Sh*^ mice was not evident until 100 dpi. This is similar to our published studies where we found that S100A8/9 deficiency resulted in reduced neutrophil lung accumulation (Scott et al., 2020), resulting in improved *Mtb* control and improved TB disease, but after 100 dpi. However, this is in contrast to the role of eosinophils in TB, as eosinophil deficiency resulted in increased *Mtb* CFU (Bohrer et al., 2021). MCs were the primary innate cell type that were defective in the lungs of *CgKitW*^*Sh*^ mice at baseline and throughout infection, and their absence was associated with better containment of *Mtb* in both lung and spleen, indicating a pathological role for MCs during chronic TB.

Our data also showed that MCs accumulated in the lungs ofC57BL/6 mice from 50 to 100 dpi, after which their numbers stabilized. This increased accumulation of MCs during chronic infection coincided with elevated infiltration of DCs and neutrophils. Importantly, we showed that enhancing MCs numbers through adoptive transfer in the lungs of WT *Mtb-infected* mice similarly promoted neutrophil accumulation and facilitated *Mtb* dissemination. This suggests that MCs are not merely a consequence of chronic inflammation, but active modulators capable of shaping immune cell dynamics early in infection. When present in sufficient numbers, MCs with their ability to promote neutrophil recruitment, can potentially influence the balance of protective versus pathological inflammation during TB. These results raise the possibility that MC accumulation during chronic infection serves to fine-tune the composition of the innate immune compartment in the lung. Whether this regulation is beneficial or detrimental to host control of *Mtb* remains to be explored. However, our data suggested that manipulating MC responses may offer a novel avenue for modulating immune dynamics in TB, particularly in the chronic phase, where inflammation must be tightly regulated to prevent tissue damage. Additionally, the reduced inflammation observed at 150 dpi is associated with decreased levels of proinflammatory cytokines and chemokines that recruit monocytes/macrophages, and T cells, and neutrophils. Overall, based on our results along with the current literature, we propose pathological roles for neutrophils and MCs, while other granulocytes, such as eosinophils, may mediate protective roles (Bohrer et al., 2021).

MCs can release cytokines and chemokines, antimicrobial peptides, and granules upon pathogen sensing and to control pathogens (Naqvi et al., 2017; Naqvi et al., 2021; Naqvi et al., 2020). In the context of *Mtb* exposure, MCs have been shown to undergo degranulation, including histamine and β-hexosaminidase (Munoz et al., 2003). Indeed, histamine-deficient mice showed decreased neutrophils, as well as proinflammatory cytokine production following *Mtb* infection (Carlos et al., 2009). Additionally, induction of degranulation following intratracheal *Mtb* infection resulted in reduced proinflammatory cytokines as well as reduced lung inflammation. Similarly, our study showed reduced DCs, neutrophils, and CD4+ and CD8+ T cell responses, in the lungs of MC-deficient *Mtb-infected* mice, along with reduced proinflammatory cytokines and lung inflammation at chronic time points. Thus, together with published work, MCs can potentially modulate neutrophils and other inflammatory mediators in high-dose models, as well as in a physiologically relevant *Mtb* infection model, leading to disease pathology. The exact mechanism by which MCs contribute to the pathology, dissemination, and promote *Mtb* infection is an area of future investigation.

At baseline, human lungs have been reported to primarily express tryptase (Poto et al., 2022). Indeed, we found that this is true for both macaque and human lungs in our study, where healthy lungs expressed MC_T_s. Additionally, we found that early granulomas and in LTBI we saw expression of MC_TC_s with a switch to more and accumulation of MCcs in late-stage granulomas. Chymase expression may modulate extracellular matrix components (FCM) such as fibronectin, leading to tissue remodeling, impacting cellular communication, and inducing cleavage for key cytokines such as IL-6, IL-13, IL-15, and IL-33, as well as TGF-p. (Pejler, 2020; Waern et al., 2013). Studies have shown that tryptase can induce proliferation of fibroblasts, epithelial cells, and smooth muscle cells, causing airway remodeling during diseased conditions (Mogren et al., 2021). Tryptase can also inactivate a large range of peptides by cleaving specific substrates, such as fibrinogen, gelatin, pro-matrix metalloproteinases (MMP), and complement factors, thus moderating inflammatory responses (Caughey, 2007). Based on our results from human and macaque lung, we hypothesize that MC_TC_S may synergize to drive responses induced by both pathways at early time points and possibly just by MC_C_s at later time points. Additional studies describing these subsets and testing their functional relevance in *in vivo* models are future steps in delineating the role of these subsets in TB. Single-cell transcriptomic analysis revealed a differential activation state between the MCs that accumulate in lungs in PTB as compared to LTBI and HC in NHPs (Esaulova et al., 2021). The MCs from PTB animals showed a closer resemblance to MC_CS_, characterized by higher IFN*γ*, metabolic activation, and chymase signatures, confirming that chymase expression is associated with disease severity. This association of chymase expression with disease severity was confirmed in an independent single-cell lung dataset (Gideon et al., 2022), where the analysis revealed similar enrichment of chymase-expressing MCcs in granulomas with higher disease burden, accompanied by similar activation of IFN-*γ* and metabolic pathways. Metabolic activation of oxidative phosphorylation, as observed here, has been associated with activated MCs, and inhibiting mitochondrial ATP production reduced MC degranulation and cytokine production (Paruchuru et al., 2022; Sharkia et al., 2017). This also suggests that MCs, which accumulate in the lungs in LTBI and HC, were not only lower in proportion but also less activated, meaning that TB-induced activation of MCs contributes to their pathogenic phenotype in disease. Our data aligned with previous observations by Garcia-Rodriguez *et al*., 2021, showing that chymase-expressing MCs accumulate in TB-induced lung lesions and may contribute to fibrotic processes surrounding granulomas (Garcia-Rodriguez et al., 2021). We also observed an expansion of MC_TC_S in the PTB group in NHPs, mirroring the phenotypic shift from M_CT_ to MC_TC_ seen in human lung sections. While Garcia-Rodriguez *et al*. suggested this shift occurs in fibrotic areas and may reflect tissue remodeling, they did not demonstrate improved lung function. Although fibrosis was not directly measured in our study, these findings support the association of chymase-positive MCs with advanced, inflammatory disease, reinforcing their potential role in TB pathogenesis.

MCs are known to gear towards a Th2 signature with increased chymase expression (Toru et al., 1998). This increase was reflected in MCs from the macaque lung, showing a high transcriptomic Th2 signature in PTB but not in clusters found in LTBI and HC. In essence, transcriptomics reflected the hyperactivated nature of the MCs in PTB, which might make them more pathological during infection. Although our *in vivo* mouse study showed a pathological role of MCs, significant secretion of Th2 cytokines was not seen, likely because the C57Bl/6 mouse model is prone towards Thl polarization (Wakeham et al., 2000). MCs can release both preformed and de novo synthesized TNF-*α*, hence helping in early bacterial clearance (Gordon & Galli, 1991). Similarly, our study showed that the MCs from HC and LTBI individuals expressed higher levels of TNF-*α* and were in a less metabolically activated state (lower OXPHOS signature). So, additional studies will help in elucidating the mechanism through which MCs mediate pathological roles.

In summary, we demonstrated that the accumulation of chymase-producing MCs in PTB is a cross-species phenomenon that contributes to increased TB pathology and loss of TB control, thereby elucidating the pathological role of MCs in the control of *Mtb* infection. By targeting MC pathways or signaling mechanisms, host-directed therapies (HDTs) hold the promise of enhancing the effectiveness of existing treatments and mitigating disease-related complications.

## Acknowledgements

This work was supported by Washington University in St. Louis; NIH grants HL105427, All 11914, AI134236, and AI123780 to S.A.K., and the Department of Microbiology, University of Chicago. We thank Ms. Lan Lu, Washington University in St. Louis, and Adlai Politi, University of Chicago, for technical support and assistance.

## Authors Contributions

S.A.K. supervised all aspects of the study. A.G. and V.T. share co-authorship. A.G., V.T. designed, performed experiments, and analyzed data. A.G., V.T., and A.A. constructed figures and analyzed data. J.R.M., N.N., Y.T., M.A., and K.S.C. curated data/contributed resources and/or data analysis. A.G., V.T., and S.A.K. wrote the original draft of the manuscript. S.A.K., A.G., V.T., A.A., and N.N. reviewed and edited the manuscript. J.Z. provided the human specimens. D.K. provided the NHP specimen, and D.K. and S.A.K. provided funding. All authors reviewed, edited, and approved the manuscript.

## Figure legends

**Figure S1:**
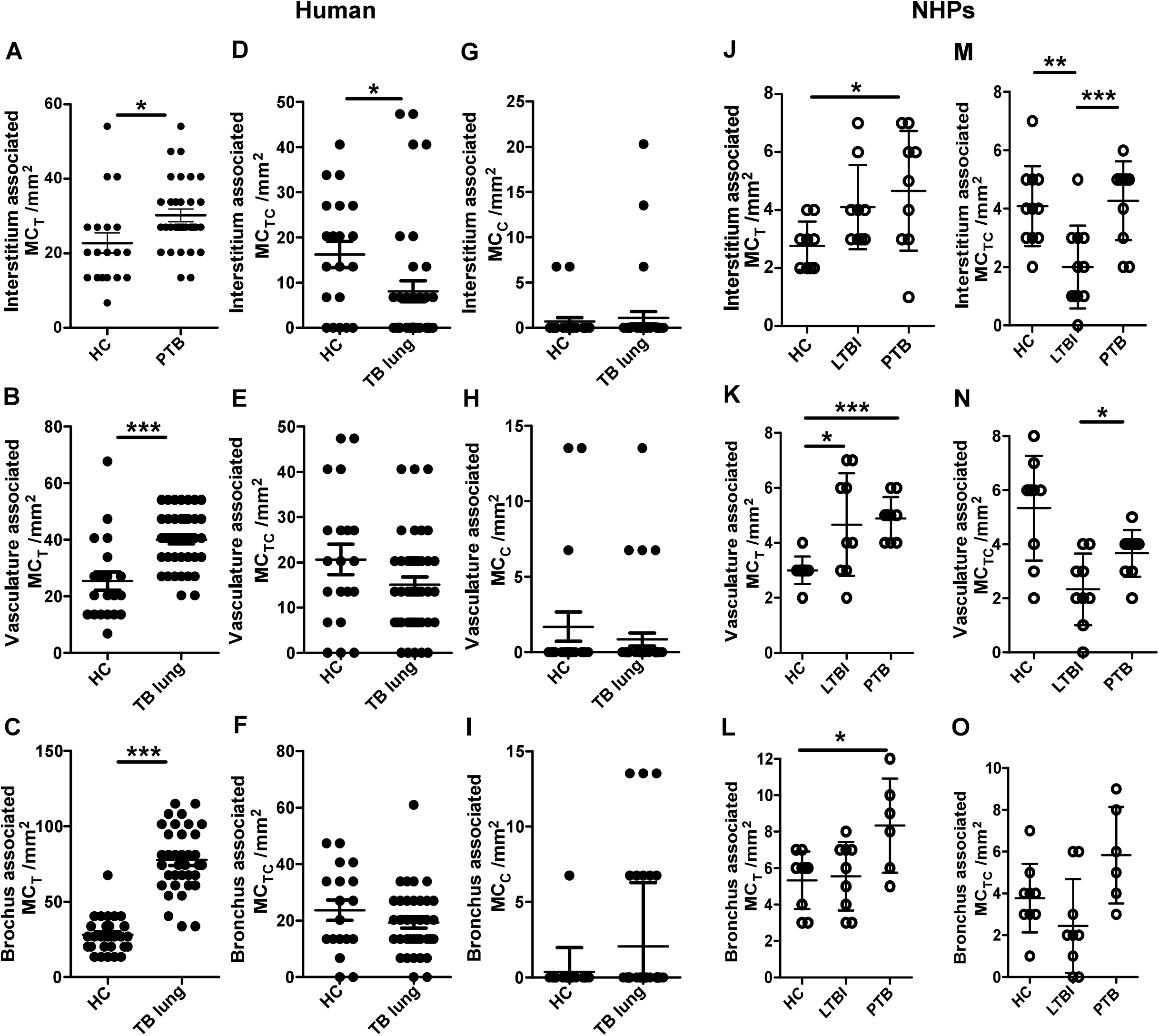
Predominance of MCTS in human and NHPs lung interstitium, blood vessels and bronchi. Healthy lung tissue and TB infected biopsies from human and NHP samples were stained for MC_T_ (green), MCc (red). Accumulation and localization of (A) MC_TS_, (B) MC_CS_. and (C) MC_TCS_ in healthy and TB infected human lung, and (D) MCTs and (E) MCTcs in LTBI and TB macaque lungs. Statistical analysis was performed using unpaired, 2-tailed Student’s t test, *** p < 0.0001, ** p < 0.001, * p< 0.05.

**Figure S2:**
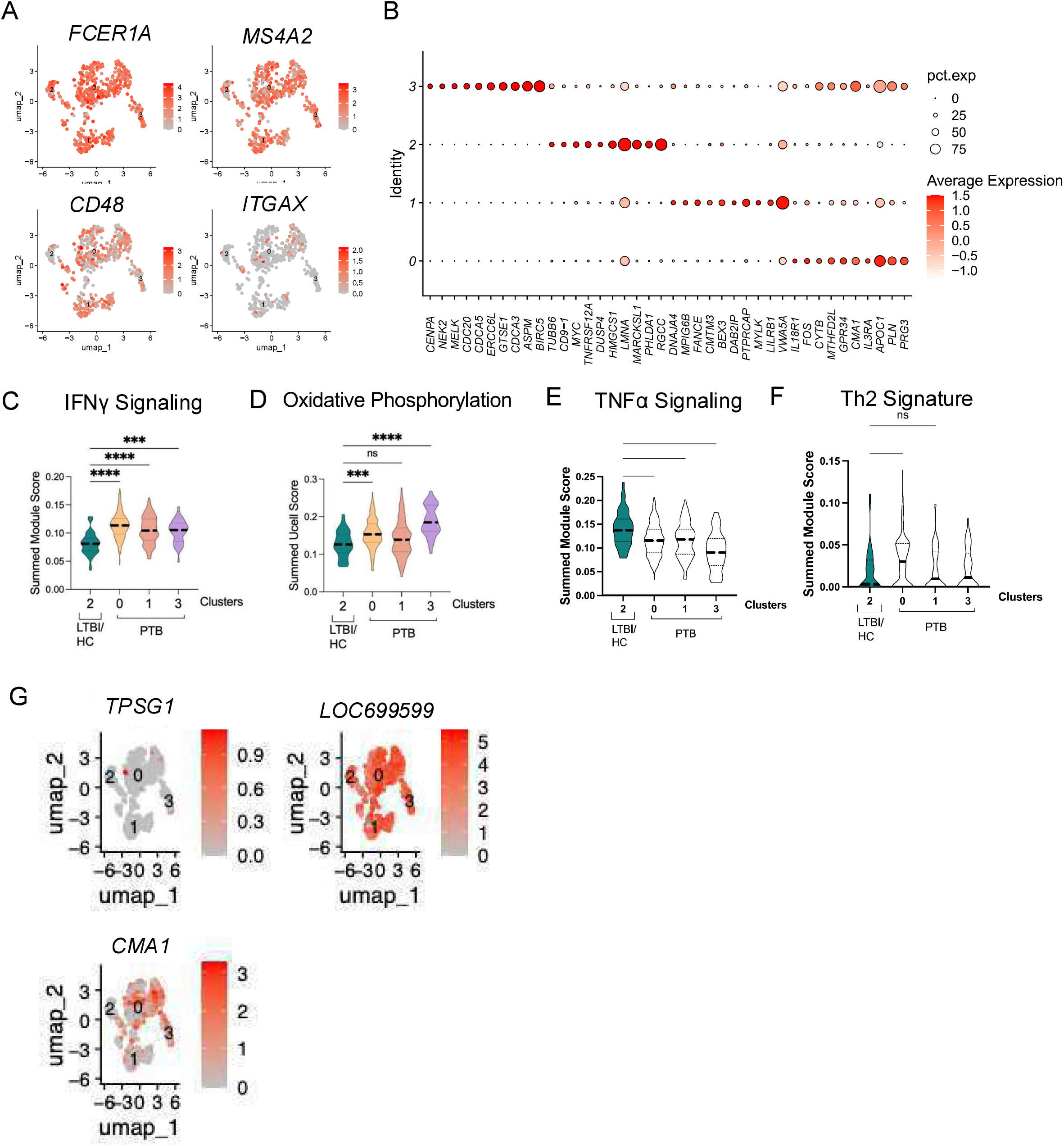
Immune cell marker expression and pathway analysis across disease conditions. Data was re-analyzed from the lungs of NHPs (GSE200151). (A) UMAP embeddings of MCs showing expression of cell surface markers: *FCER1A, MS4A2, CD48*, and *ITGAX* across identified clusters. (B) Dot plot showing the expression levels of select genes across clusters, where the dot size represents the percentage of cells expressing each gene, and the color intensity represents the average expression level. (C) Violin plot of summed module scores for IFNy signaling, Oxidative Phosphorylation (D), TNF-*α* signaling (E), Th2 signature (F) pathway across different disease conditions: LTBI/HC (green), PTB (pink), and the respective clusters. (G) UMAP embeddings show the expression of key MC markers, including *TPSG1, LOC699599*, and *CMA1* across identified clusters, with color intensity indicating expression levels. Statistical significance was assessed using Kruskal-Wallis tests with Dunn’s multiple comparison correction (**p < 0.01, ***p < 0.001, ****p < 0.0001, ns: not significant).

**Figure S3:**
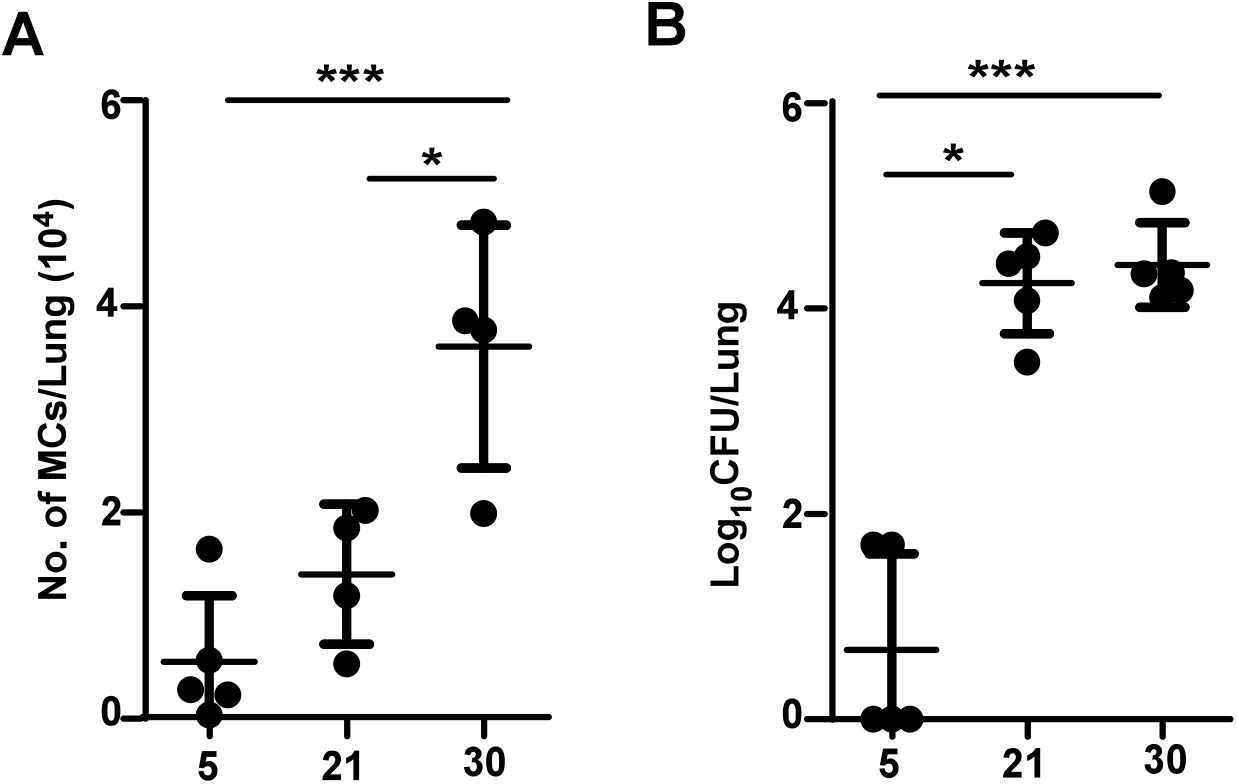
MCs appear at early *Mtb* infection. C57BL/6 mice were infected with a low aerosol dose (-l00CFU) of *Mtb* HN878 and mice were sacrificed at 5, 21 and 30 dpi. (A) The number of MCs in the lungs was enumerated in Mtb-infected mice. (B) Bacterial burden was assessed by plating. Data points represent the mean± SD *(n* = 4-5 per time point). Statistical analysis was performed using an unpaired, 2-tailed Student’s t test between time points, *** p < 0.0001, ** p < 0.001, * p< 0.05.

**Figure S4:**
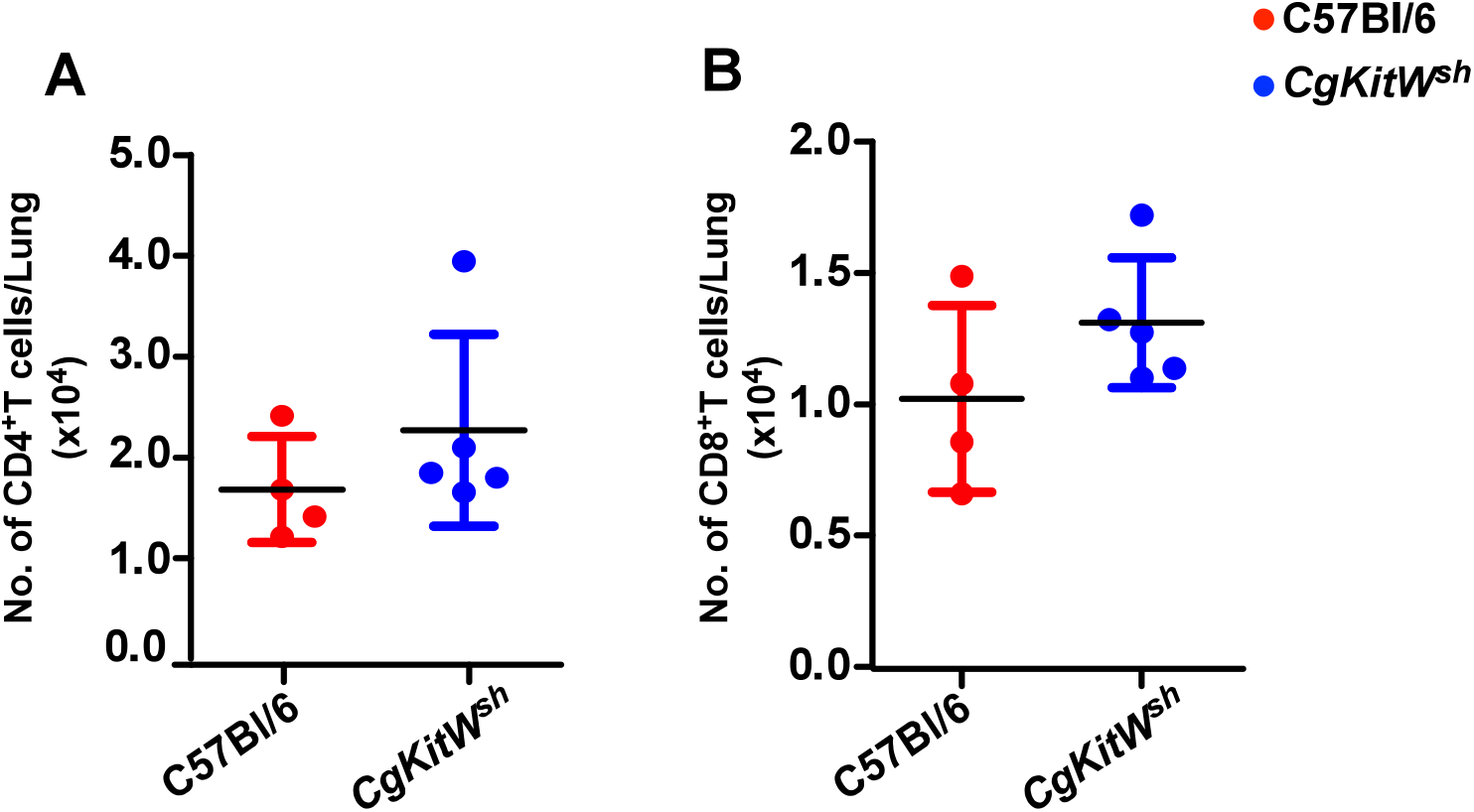
MC-deficient mice have no baseline differences in T cell numbers. Six weeks C57BL/6 mice and *CgKitW*^*Sh*^ mice were sacrificed to enumerate baseline numbers of (A) CD4+T cells, and of (A) CD4 ^+^ T cells, and (B) CD8^+^ T cells. Data points represent the mean ± SD (*n* = 4-5 per time point).

**Figure S5:**
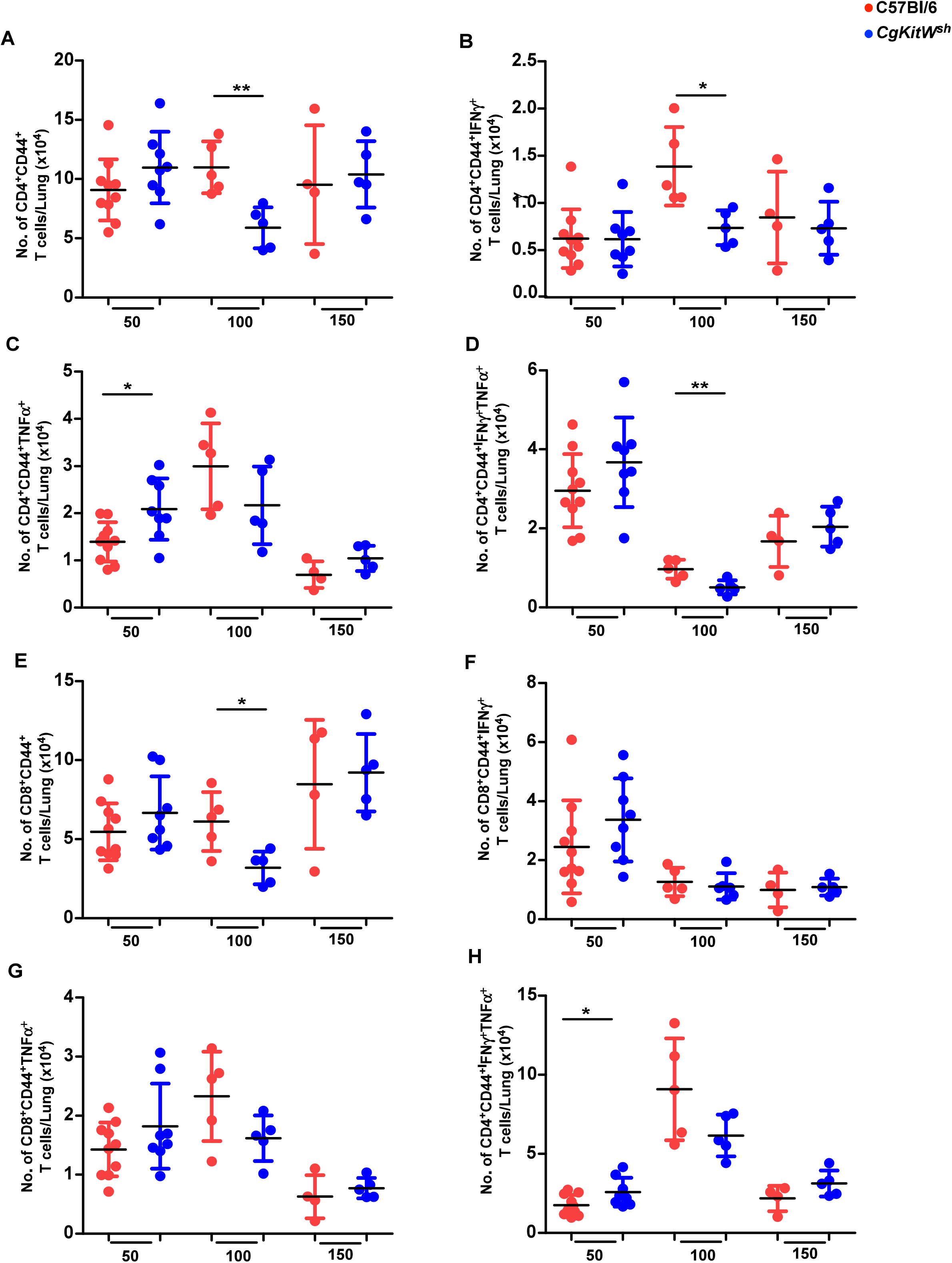
MC-deficient mice have reduced numbers of activated CD4+ and cos+ T cells in the lung. C57BL/6 and CgKitW^sh^ mice were infected with a low aerosol dose (∼l00CFU) of *Mtb* HNS7S and mice were sacrificed at 50, 100 and 150 dpi. Number of(A) CD4^+^CD44^+^T cells, (B) CD4^+^CD44^+^IFN*γ*^+^ T cells, (C) CD4^+^CD44^+^TNF-*α*^+^T cells (D) CD4^+^CD44^+^IFN*γ*^+^ TNF--*α*^+^T cells, (E) CD8^+^CD44^+^T cells, (F) CD8+ CD44^+^IFN*γ*^+^T cells, (G) CD8^+^ CD44^+^ TNF-*α*^+^T cells, and (H) CD8^+^CD44^+^IFN*γ*^+^ TNF-*α*^+^T cells in the lungs of *Mtb* infected mice. Data points represent the mean± SD of 1 of 2 individual experiments *(n* = 4-10 per time point per group). Statistical analysis was performed using unpaired, 2-tailed Student’s t test between C57BL/6 and CgKitW^sh^ mice,*** p < 0.0001, ** p < 0.001, * p< 0.05. Outliers were removed from the subsets using Grubb’s outlier test.

**Figure S6:**
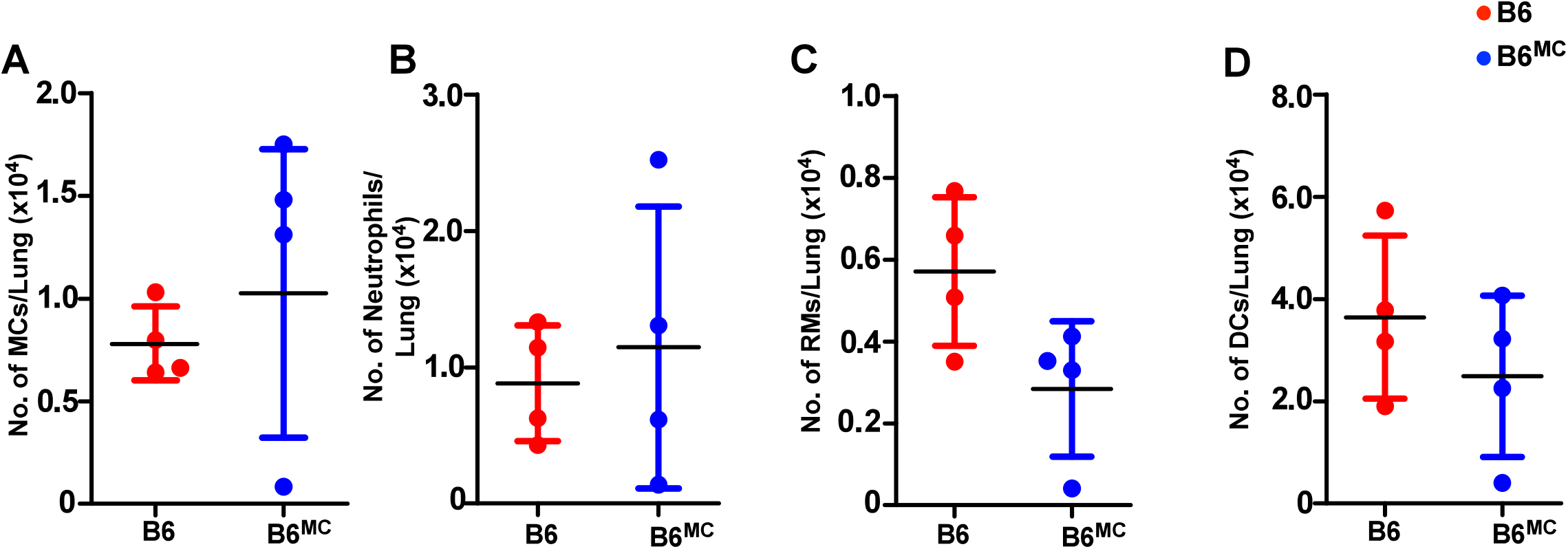
Lung myeloid cell accumulation did not vary in MC-transferred WT *Mtb* infected mice. Bone marrow derived in vitro cultured MCs (n=50,000 cells per mouse) were adoptively transferred into the lung airways of WT mice 7 days before infecting with a low aerosol dose (∼ l00CFU) of *Mtb* HNS78. MCs were replenished in these mice at 15 dpi, and mice were sacrificed at 30 dpi. Numbers of (A) MCs, (B) neutrophils, and (C) RMs were enumerated in the lungs of *Mtb-infected* mice. Data points represent the mean ± SD of 1 of 2 individual the lungs of *Mtb*-infected mice. Data points represent the mean ± SD of 1 of 2 individual experiments (*n* = 4 per group).

